# Single molecule counting detects low-copy glycine receptors in hippocampal and striatal synapses

**DOI:** 10.1101/2025.01.07.631736

**Authors:** Serena Camuso, Yana Vella, Souad Youjil Abadi, Clémence Mille, Bert Brône, Christian G Specht

**Affiliations:** NeuroBicêtre, Inserm U1195, Université Paris-Saclay, 94276 Le Kremlin-Bicêtre, France; UHasselt, Neurophysiology Laboratory, BIOMED research institute, 3500 Hasselt, Belgium

## Abstract

Glycine receptors (GlyRs) are heteropentameric chloride channels that mediate fast inhibitory neurotransmission in the brainstem and spinal cord, where they regulate motor and sensory processes. GlyRs are clustered in the post-synaptic membrane by strong interactions of the β subunit with the scaffold protein gephyrin. Even though GlyRβ mRNA is highly expressed throughout the brain, the existence of synaptic GlyRs remains controversial as there is little conclusive evidence using conventional fluorescence microscopy and electrophysiological recordings. Here we exploit the high sensitivity and spatial resolution of single molecule localisation microscopy (SMLM) to investigate the presence of GlyRs at inhibitory synapses in the brain, focusing on several areas of the telencephalon. Making use of a knock-in mouse model expressing endogenous mEos4b-tagged GlyRβ, we identified few GlyRs in sub-regions of the hippocampus. Dual-colour SMLM revealed that these sparse receptors are integrated within the post-synaptic gephyrin domain, pointing to a possible role in maintaining the structural integrity of inhibitory synapses. In contrast, we found functionally relevant numbers of synaptic GlyRs at inhibitory synapses in the ventral striatum. Our results highlight the strength of SMLM to detect few and sparsely distributed synaptic molecules in complex samples and to analyse their organisation with high spatial precision.

## Introduction

Inhibitory neurotransmission in the central nervous system (CNS) is largely mediated by glycine receptors (GlyRs) and γ-aminobutyric acid type A receptors (GABAARs) (Alvarez, 2017, Kasaragod & Schindelin, 2018). Both classes of receptor are widely expressed throughout the brain and the spinal cord, however, they have specific regional expression patterns. GlyRs are particularly abundant in the spinal cord and brain stem, where they play an important role in the processing of sensory and motor information as well as the modulation of pain responses (Alvarez, 2017, Fenech et al., 2024). Although the expression of GlyRs is much lower in the brain (Maynard et al., 2021, Zeilhofer et al., 2005), glycinergic transmission is known to be involved in reward signalling and possibly pain associated responses (Adermark et al., 2011, Devoght et al., 2023, Fenech et al., 2024, Muñoz et al., 2018, San Martin et al., 2020).

Both GABA_A_Rs and GlyRs are cys-loop pentameric chloride channels that are composed of different combinations of subunits. Each subunit contains a large N-terminal extracellular domain (ECD), four transmembrane domains (TM1-4), as well as a flexible intracellular domain (ICD) between TM3 and TM4 (Kasaragod & Schindelin, 2018). Pentameric GlyRs are assembled from five different subunits, α1-α4 and β. Homopentameric GlyRs composed only of α subunits are mostly found in the extra-synaptic plasma membrane. In contrast, heteropentameric receptors containing both α and β subunits accumulate at post-synaptic sites, due to a direct interaction between the ICD of the β subunits and gephyrin, the main scaffold protein at inhibitory synapses (Alvarez, 2017).

*In situ* hybridisation studies have shown that the mRNA of the GlyRα1 and β subunits is highly expressed in spinal cord neurons (Ceder et al., 2024, Malosio et al., 1991). The GlyRβ transcript is also expressed in most brain regions, including olfactory bulb, cerebral cortex, hippocampus, and striatum (Ceder et al., 2024, Fujita et al., 1991, Malosio et al., 1991). Surprisingly, expression of GlyRβ protein appears to be exceedingly low in the telencephalon. For example, GlyR labelling was only detected at a few synapses in the hippocampus, mainly in the pyramidal layer (Danglot et al., 2004, Maynard et al., 2021, Weltzien et al., 2012). In line with this, electrophysiological measurements have failed to detect synaptic GlyR currents in the hippocampus (Chattipakorn & McMahon, 2002, Mori et al., 2002, Song et al., 2006), with only a single study reporting evoked IPSCs in mouse hippocampal CA1pyramidal cells (Muller et al., 2013).

The mRNA levels of the GlyR α subunits are generally low across the brain (Ceder et al., 2024, Malosio et al., 1991). Yet, surface expression of homopentameric GlyRs was demonstrated by electrophysiological recordings of glycine-induced currents in several regions including the hippocampus and dorsal striatum (Chattipakorn & McMahon, 2002, Comhair et al., 2018, Molchanova et al., 2017, Mori et al., 2002, Song et al., 2006). These extrasynaptic receptors are thought to play a role in the tonic inhibition of central neurons (McCracken et al., 2017, Mori et al., 2002, Song et al., 2006). This situation appears to be different in the ventral striatum (*nucleus accumbens*), where glycinergic mIPSCs likely corresponding to heteropentameric GlyRs have been reported (Muñoz et al., 2018), in addition to homopentameric receptors (McCracken et al., 2017, Molander & Söderpalm, 2005, Muñoz et al., 2020).

To explore the presence of GlyRs at synapses in the brain, we made use of the high spatial resolution and extraordinary sensitivity of single molecule localisation microscopy (SMLM), which enabled us to detect individual GlyR complexes in different regions of the telencephalon, including the hippocampal formation and striatum. Using a knock-in (KI) mouse model that expresses endogenous mEos4b-tagged GlyRβ subunits (Maynard et al., 2021), we identified low copy numbers of GlyRs at inhibitory synapses. Dual-colour SMLM further demonstrated the integration of these GlyRs within the post-synaptic gephyrin scaffold, suggesting that they are important for the assembly or maintenance of inhibitory synaptic structures in the brain.

## Results

### Identification of low-copy synaptic GlyRs in mouse hippocampus

The distribution of glycinergic synapses is mainly confined to the spinal cord and brain stem (Alvarez, 2017). While the presence of GlyRβ subunit mRNAs is well documented in different brain areas (Ceder et al., 2024, Fujita et al., 1991, Malosio et al., 1991); see also Supplementary Fig. S1), little is known about GlyRβ protein expression in the brain (Danglot et al., 2004, Maynard et al., 2021). Here, we exploited the high sensitivity and spatial resolution of SMLM to probe the presence of GlyRβ subunits at synapses in the brain, focusing on the hippocampus.

Since no reliable antibodies against the β-subunit of the GlyR are available, we used a knock-in mouse model expressing endogenous mEos4b-tagged GlyRβ subunits (Maynard et al., 2021, Wiessler et al., 2024). No obvious functional, behavioural or ultrastructural phenotypes have been reported in homozygous and heterozygous animals expressing mEos4b-GlyRβ. Thin cryostat coronal sections (10 µm) were cut from homozygous *Glrb*^eos/eos^ mouse brain and labelled with NeuN antibody and with Sylite, a small peptide probe against the inhibitory synaptic scaffold protein gephyrin (Khayenko et al., 2022). The analyses were conducted in the molecular layer of the dentate gyrus (DG), where the dendrites of granule cells receive synaptic inputs from the entorhinal cortex, and in the *stratum radiatum* of the CA3 and CA1 regions, where the apical dendrites of pyramidal cells make contact with Mossy fibers and Schaffer collaterals, respectively (Supplementary Fig. S1C). The different hippocampal sub-regions were identified using the neuronal marker NeuN in the green channel (not shown). Reference images of Sylite were taken in the far-red channel, followed by SMLM recordings of the green-to-red photoconvertible fluorescent protein mEos4b attached to the GlyRβ subunit (Fig. 1A).

**Figure 1.**
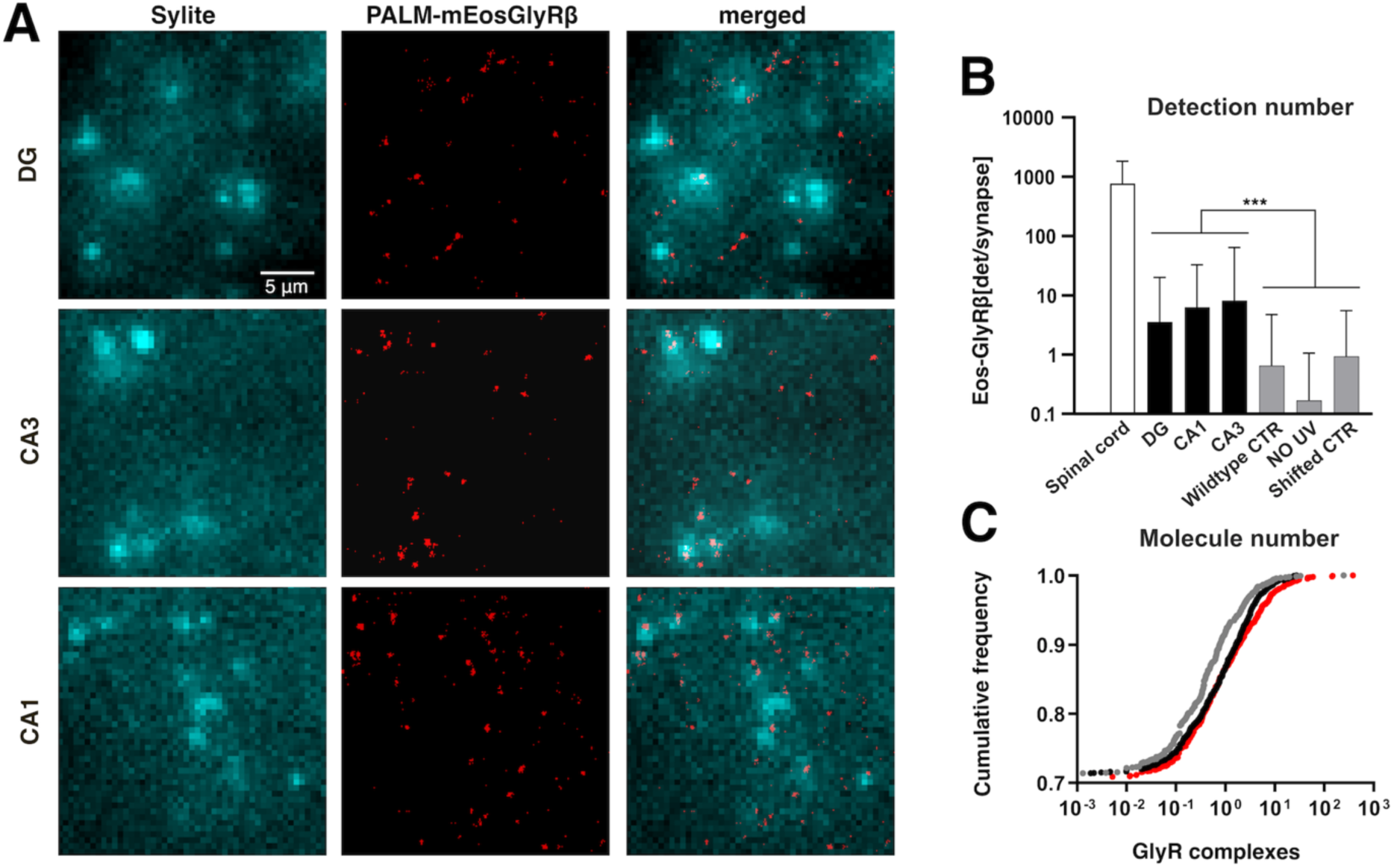
SMLM of mEos4b-GlyRβ subunits in the hippocampus. **(A)** Single molecule detections of mEos4b-GlyRβ (red SMLM pointillist image) in the DG, CA1 and CA3 regions of the hippocampus in 10 µm cryostat sections of the knock-in mouse line *Glrb*^eos/eos^ at postnatal day 40 (red pointillist images). Inhibitory synapses were identified in epifluorescence images using the gephyrin marker Sylite (cyan). Scale bar 5 µm. **(B)** Mean number of mEos4b-GlyRβ detections per synaptic gephyrin cluster in spinal cord (n = 9733 synaptic clusters from 10 fields of view) and brain slices (n > 5000 clusters from 9 fields of view for each region) from N = 3 independent experiments corresponding to three *Glrb*^eos/eos^ mice. Recordings were also made in the CA3 of wildtype mice not expressing endogenous mEos4b-GlyRβ (negative control; n = 3637 clusters, 3 fields of view, 2 *Glrb*^WT/WT^ mice) and in *Glrb*^eos/eos^ hippocampal slices without photoconversion of mEos4b (no UV, n = 3301 clusters, 9 fields of view, 3 animals). A pixel shift control with flipped images was done with the same dataset (*Glrb*^eos/eos^, CA3; n = 3301 clusters, 9 fields of view, 3 animals). Data are shown as mean ± SD. Levels of significance were determined using a Kruskal-Wallis test with Dunn’s multiple comparison test: *** *p* < 0.0001. **(C)** Cumulative distribution representing the estimated copy number of mEos4b-GlyRβ complexes per synapse in DG (grey line), CA1 (black line) and CA3 (red line) regions. Copy numbers were background-corrected by subtracting the value obtained for the negative control in wildtype slices.

We observed very few single molecule detections during SMLM, indicating exceedingly low mEos4b-GlyRβ expression in the hippocampus. SMLM super-resolution images were reconstructed and analysed using Icy software to quantify the number of mEos4b detections per gephyrin cluster (Fig. 1B). We counted on average between 3 to 10 detections per synapse in all hippocampal sub-regions. This is about two orders of magnitude lower than in the spinal cord (*p* < 0.0001, non-parametric Kruskal-Wallis one-way ANOVA with Dunn’s multiple comparison test, KW test), where we counted almost 1000 detections of mEos4b-GlyRβ per synapse (Fig. 1B, Supplementary Fig. S2). Aside from the low detection numbers in the hippocampus, we did not find obvious differences between the sub-regions, with a significant difference only between the DG and CA1 (*p =* 0.002, KW test).

To ascertain that what we saw in the hippocampus were indeed mEos4b detections and not imaging artefacts, we performed control experiments in hippocampal slices from wildtype mice not expressing mEos4b-tagged GlyRβ subunits. As expected, the mean number of detections at CA3 synapses was significantly lower in the negative control than in the *Glrb*^eos/eos^ slices (Fig. 1B; *p* < 0.0001, KW test). Similarly, we recorded SMLM movies in *Glrb*^eos/eos^ slices without 405 nm illumination. In the absence of UV, the mEos4b fluorescent protein is not converted into its red form, making this a stringent internal control. The number of detections per gephyrin cluster was again much lower than in the recordings with 405 nm laser illumination (Fig. 1B; *p* < 0.0001, KW). These data confirm that the non-specific background of detections, mainly outside of synapses, is sufficiently low to accurately identify and quantify endogenous mEos4b-GlyRβ subunits in the hippocampus of our knock-in mouse model. To exclude the possibility that the co-localisation between the mEos4b-GlyRβ detections and gephyrin was due to a random overlap, we did a pixel shift analysis with two-colour images of the CA3 region, in which the Sylite channel was horizontally flipped relative to the SMLM image of the mEos4b detections. This transformation led to a significant reduction in the number of mEos4b-GlyRβ detections at gephyrin clusters compared to the original data (Fig. 1B; *p* < 0.0001, KW), confirming that the localisation of mEos4b-GlyRβ at inhibitory synapses is not down to chance.

We also estimated the absolute number of GlyRs per synapse in the hippocampus. The number of mEos4b detections was converted into copy numbers by dividing the detections at synapses by the average number of detections of individual mEos4b-GlyRβ containing receptors. This value was measured across CA3 slices, both at synapses as well as in the extrasynaptic region (see Methods). The advantage of this counting strategy is that it is independent of the stoichiometry of heteropentameric GlyRs that is still under debate (e.g. (Durisic et al., 2012, Grudzinska et al., 2005, Patrizio et al., 2017, Zhu & Gouaux, 2021)). The obtained copy numbers are reported as background-subtracted values (Table 1). According to our quantification GlyRs are present at about a quarter of hippocampal synapses, with copy numbers most often in the single digits (Fig. 1C, Table 1). Only few synapses contain 10 or more GlyRs, and as many as 75% of hippocampal synapses do not contain any GlyRs at all. No obvious differences were noticed in the copy number of GlyRs between sub-regions. Taken together, our findings demonstrate the existence of very few heteropentameric GlyR complexes at inhibitory synapses in the hippocampus. With an average of approximately one GlyR per synapse, this is about a hundredfold lower than in the spinal cord.

**Table 1.** Estimated copy numbers of mEos4b-GlyRβ containing heteropentameric GlyR complexes at inhibitory synapses in different regions of the CNS of *Glrb*^eos/eos^ KI mice (background-corrected, see Methods).

| CNS region | GlyR copy number<br>(mean ± SEM) | range of GlyR copies<br>(5-95 percentile) | fraction of GlyR positive<br>synapses ($\geq 0.5$ copies) |
| --- | --- | --- | --- |
| Spinal cord | 120 ± 5 | 0 – 470 | 0.89 |
| CA1 | 0.34 ± 0.05 | 0 – 3 | 0.18 |
| CA3 | 1.11 ± 0.30 | 0 – 5 | 0.18 |
| DG | 0.35 ± 0.18 | 0 – 2 | 0.13 |
| Dorsal striatum | 3.00 ± 0.18 | 0 – 15 | 0.43 |
| Ventral striatum | 26.10 ± 1.48 | 0 – 112 | 0.73 |

### Sub-synaptic distribution of GlyRs at hippocampal synapses

At glycinergic synapses in the spinal cord the receptors are anchored in the post-synaptic membrane by gephyrin, a scaffold protein that binds with high affinity to the GlyRβ subunit, promoting the synaptic localisation of GlyRs (e.g. (Kasaragod & Schindelin, 2018, Kostrz et al., 2024, Maynard et al., 2021)). The nanoscale organisation of inhibitory synapses in the hippocampus containing low-copy GlyRs has not yet been studied in detail (Danglot et al., 2004). To obtain information about the distribution of GlyRs within the synaptic structure, we carried out dual-colour SMLM of GlyRβ and gephyrin in brain slices of adult *Glrb*^eos/eos^ mice using stochastic optical reconstruction microscopy (dSTORM) with organic fluorophores in a reducing buffer (Yang & Specht, 2020). Compared to SMLM with fluorescent proteins, dSTORM provides higher photon yields, resulting in a better localisation precision (∼12 nm in our recordings, see Methods), which is critical for resolving fine structural details. For these experiments, we focused on the CA3 region of the hippocampus, where the presence of GlyRs at inhibitory synapses was demonstrated before (Fig. 1). Due to their low number, we did not expect to see any particular organisation of the receptors. Instead, the aim was to determine whether GlyRs are actually intrinsic components of the postsynaptic domain of inhibitory hippocampal synapses.

Endogenous mEos4b-GlyRβ subunits in tissue slices of *Glrb*^eos/eos^ KI animals were labelled with anti-mEos-AF647 nanobodies (NanoTag). The specificity of the nanobody was tested in spinal cord slices that were also labelled with gephyrin antibody (mAb7a, Synaptic Systems) and a CF568-conjugated secondary antibody. Wide-field fluorescence images showed extensive co-localisation of all three channels (mEos4b, anti-gephyrin-CF568, anti-mEos-AF647) in *Glrb*^eos/eos^ slices; the non-specific background of the nanobody in wildtype animals was very low (Supplementary Fig. S3).

For dual-colour SMLM, brain slices from *Glrb*^eos/eos^ mice were labelled with anti-mEos-AF647 nanobody as well as with anti-gephyrin and CF680-conjugated anti-mouse secondary antibodies (Fig. 2A). The emitted light from the two far-red dyes was separated by spectral demixing, using a SAFe 360 nanoscope (Abbelight) equipped with a dichroic mirror at 700 nm and two sCMOS cameras for detection. In this imaging modality, the single molecule detections recorded simultaneously on the two cameras are attributed to one or the other far-red fluorophore based on their specific intensity ratio (see Methods; Supplementary Fig. S4A-E). To further test the specificity of the nanobody labelling and the demixing procedure, we performed experiments in *Glrb*^eos/eos^ slices in which the nanobody was omitted and only gephyrin was labelled (CF680). Under these conditions, spectral demixing produced a single peak of intensity ratios corresponding to the CF680 fluorophore, with only few wrongly attributed detections in the AF647 channel (Supplementary Fig. S4F,G).

**Figure 2.**
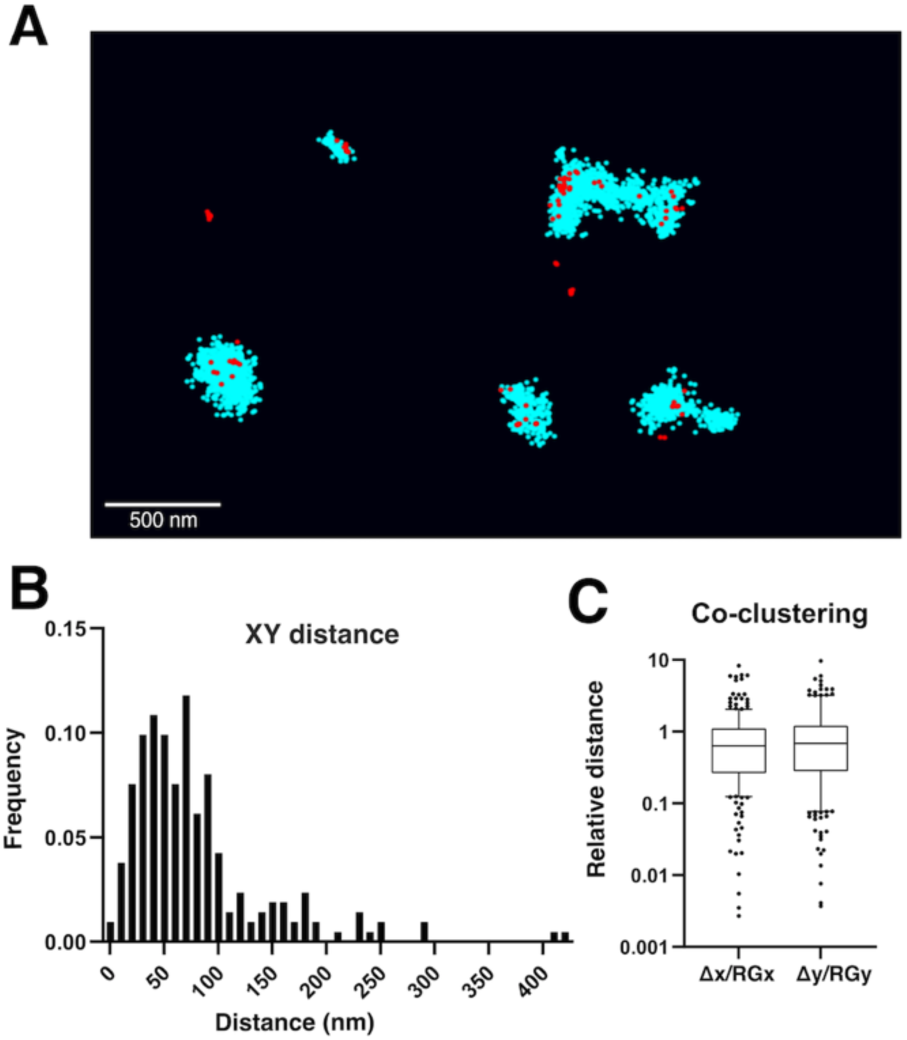
Dual-colour SMLM of GlyR and gephyrin at inhibitory hippocampal synapses. **(A)** Dual-colour SMLM using spectral demixing of endogenous mEos4b-GlyRβ labelled with anti-mEos-AF647 nanobody (red, NanoTag), and mouse anti-gephyrin (mAb7a, Synaptic Systems) and CF680-conjugated secondary anti-mouse antibodies (cyan) in hippocampal slices of the *Glrb*^eos/eos^ knock-in mouse line at postnatal day 40. Scale: 500 nm. **(B**) Euclidean distance between the centre of mass (CM) of the anti-mEos-AF647 nanobody (GlyRβ) and gephyrin (mAb7a-CF680) detections of corresponding clusters. **(C)** Distance of the CM of the anti-mEos-AF647detections from the CM of gephyrin relative to the radius of gyration (RG) of the gephyrin cluster along the x and y axes (Δx/RGx, Δy/RGy). N = 2 independent experiments corresponding with 2 *Glrb*^eos/eos^ animals.

In line with our SMLM experiments with the photoconvertible fluorescent protein mEos4b in brain slices (Fig. 1), we observed sparse single molecule detections of anti-mEos-AF647 nanobody that generally co-localised with dense clusters of gephyrin detections, confirming the presence of mEos4b-GlyRβ at inhibitory synapses in the hippocampus (Fig. 2A). The two-colour SMLM pointillist images were analysed using the DBSCAN clustering tool in the NEO software to measure the distance between mEos4b-GlyRβ and the corresponding gephyrin clusters. According to our measurements, the Euclidean distance between the centre of mass (CM) of GlyRβ and gephyrin was 80 ± 69 nm (mean ± SD, n = 214 pairs of clusters), pointing to a close spatial relationship at inhibitory synapses (Fig. 2B). We also calculated the relative distance of the GlyRβ detections from the centre of mass of gephyrin, compared to the mean distance of the gephyrin detections from their own centre of mass as defined by the radius of gyration (RG) of the gephyrin clusters. This ratio was below one for most clusters, indicating that the GlyRs are well integrated within the postsynaptic gephyrin domains at CA3 synapses (Fig. 2C).

### Differential expression of endogenous mEos4b-GlyRβ in dorsal and ventral striatum

Several reports have described the presence of glycinergic currents in the striatum (Molchanova et al., 2017, Muñoz et al., 2018). However, the localisation of GlyRs at synapses in this brain area remains controversial. Homopentameric GlyRα2 complexes are thought to mediate tonic inhibition in the dorsal striatum (Devoght et al., 2023, Molchanova et al., 2017), yet glycinergic miniature inhibitory postsynaptic currents (mIPSCs) were detected in the *nucleus accumbens*, i.e. part of the ventral striatum (Muñoz et al., 2018). To resolve this issue, we quantified the GlyRβ-containing synaptic receptors in the striatum of *Glrb*^eos/eos^ animals using SMLM.

The number of detections of mEos4b-GlyRβ per gephyrin cluster, as well as the resulting copy number of GlyRs at synapses were measured in the dorsal and in the ventral striatum in *Glrb*^eos/eos^ mice at postnatal day 40. Coronal cryo-sections of 10 µm thickness were immunolabeled for NeuN to distinguish the two sub-regions of the striatum. Gephyrin was labelled with the Sylite probe in order to identify inhibitory synapses (Fig. 3A). The average number of mEos4b-GlyRβ detections per gephyrin cluster was significantly higher in the ventral striatum compared to the dorsal striatum (136 ± 5 versus 25 ± 1 detections per synapse, mean ± SEM, n = 3309 and 2352 synapses, respectively; *p* < 0.0001, non-parametric Mann-Whitney test, MW test; Fig. 3B). Conversion of the detection numbers into receptor copy numbers revealed that about 40% of synapses in the ventral striatum contain at least 10 heteropentameric GlyR complexes, many more than in the dorsal striatum (*p* < 0.0001, KW test; Fig. 3C, Table 1). As before, copy numbers are given as background-corrected values (see Methods). The same quantification was done in the spinal cord, where we found significantly higher copy numbers of receptor complexes containing mEos4b-GlyRβ (*p* < 0.0001 against both striatal regions, KW). With an average of 120 ± 5 receptor complexes per synaptic cluster (mean ± SEM, n = 4006 synapses, Table 1), these values are close to previous estimates (Maynard et al., 2021).

**Figure 3.**
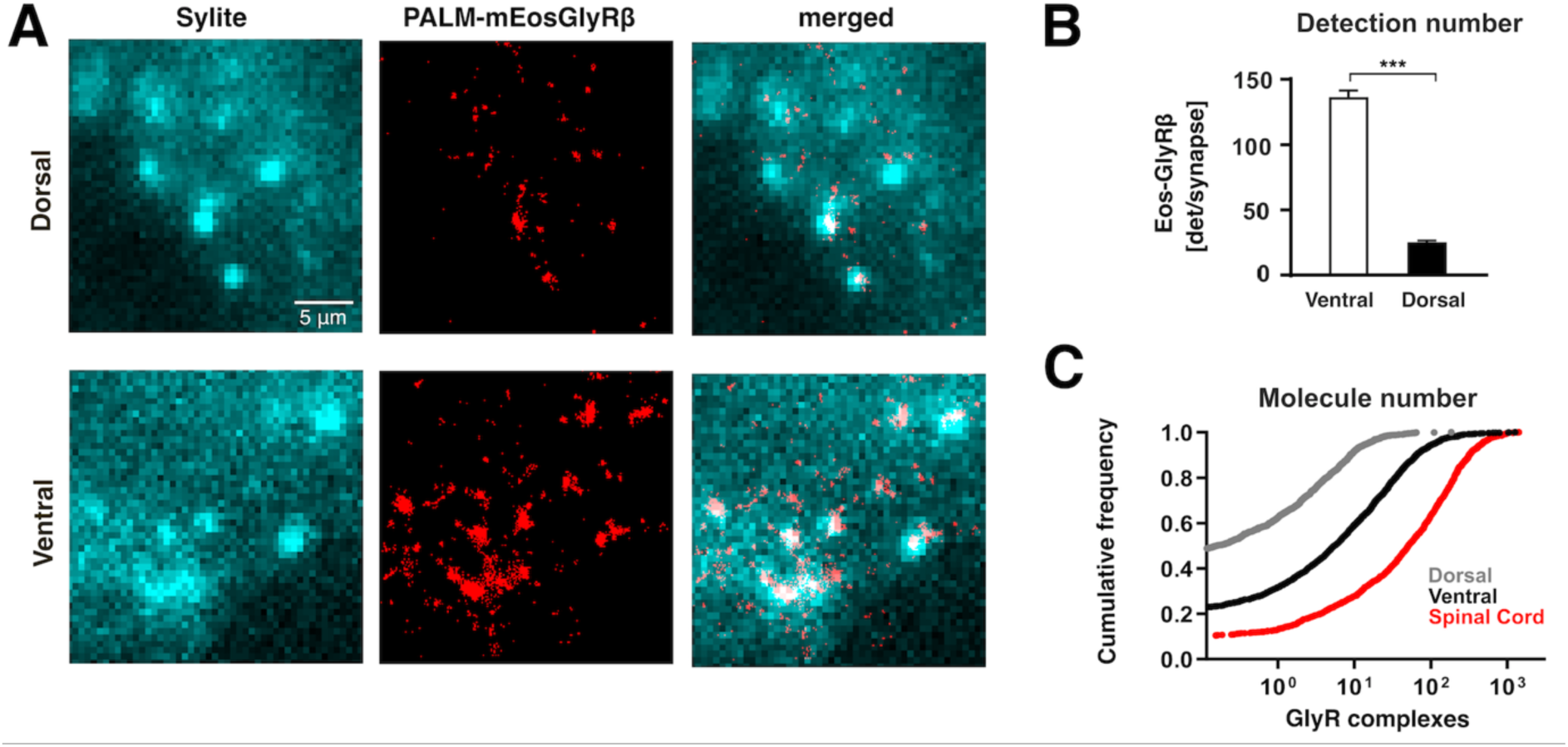
Quantitative SMLM of endogenous GlyRs at synapses in the striatum. **(A)** Super-resolution imaging of mEos4b-GlyRβ subunits (in red) in the dorsal and ventral striatum of *Glrb*^eos/eos^ knock-in mice. Cryostat slices were labelled with the gephyrin marker Sylite (cyan). Scale: 5 µm. **(B)** Single molecule detection numbers of mEos4b-GlyRβ per gephyrin cluster (n = 3309 and 2352 clusters for ventral and dorsal striatum, respectively, from 9 fields of view per sub-region and N = 3 independent experiments from three animals; mean ± SEM; two-tailed Mann-Whitney test: *** *p* < 0.0001). **(C)** Cumulative distribution of the estimated number of mEos4b-GlyRβ containing receptor complexes per synapse in the dorsal striatum (grey line), ventral striatum (black line) and spinal cord (red line). Copy numbers are background corrected (see Table 1).

Our findings show that the numbers of mEos4b-GlyRβ subunits are much higher at synapses in the striatum compared to synapses in the hippocampal formation, and that their distribution in the striatum is area-specific. This observation is supported by electrophysiological recordings of glycinergic synaptic currents in the *nucleus accumbens* (Muñoz et al., 2018) as opposed to mainly extrasynaptic tonic currents in the dorsal striatum (Molchanova et al., 2017).

### Miniature synaptic currents in ventral striatum recorded with whole cell patch-clamp

Since the published recordings of glycinergic currents in the striatum were performed under different conditions, i.e. different striatal sub-regions, mouse strains and ages (Molchanova et al., 2017, Muñoz et al., 2018), we wanted to verify the area-specific functional differences in a single experimental setting. We pharmacologically isolated glycinergic mIPSCs and determined their frequency and amplitudes in brain slices of both ventral and dorsal striatum of C57BL/6J mice at postnatal day 35 to 41 using whole cell patch clamp (Fig. 4). mIPSCs are considered a measure of functional synapses because they are generated by the spontaneous release of synaptic vesicles from presynaptic terminals. Our data confirm the presence of glycinergic mIPSCs in medium spiny neurons (MSNs) in the ventral striatum and the near complete absence of glycinergic mIPSCs in dorsal striatal MSNs (*p* < 0.05, MW test; Fig. 4D). With an amplitude of 17.2 ± 2.9 pA (mean ± SEM, n = 5 cells from N = 3 animals; Fig. 4C) and a frequency of 0.12 ± 0.03 Hz (Fig. 4D), the mIPSCs recordings in the ventral striatum are similar to the results of an earlier study of the *nucleus accumbens* (Muñoz et al., 2018). Our electrophysiological data also match the differences in synaptic GlyR copy numbers obtained by quantitative SMLM (Fig. 3C).

**Figure 4.**
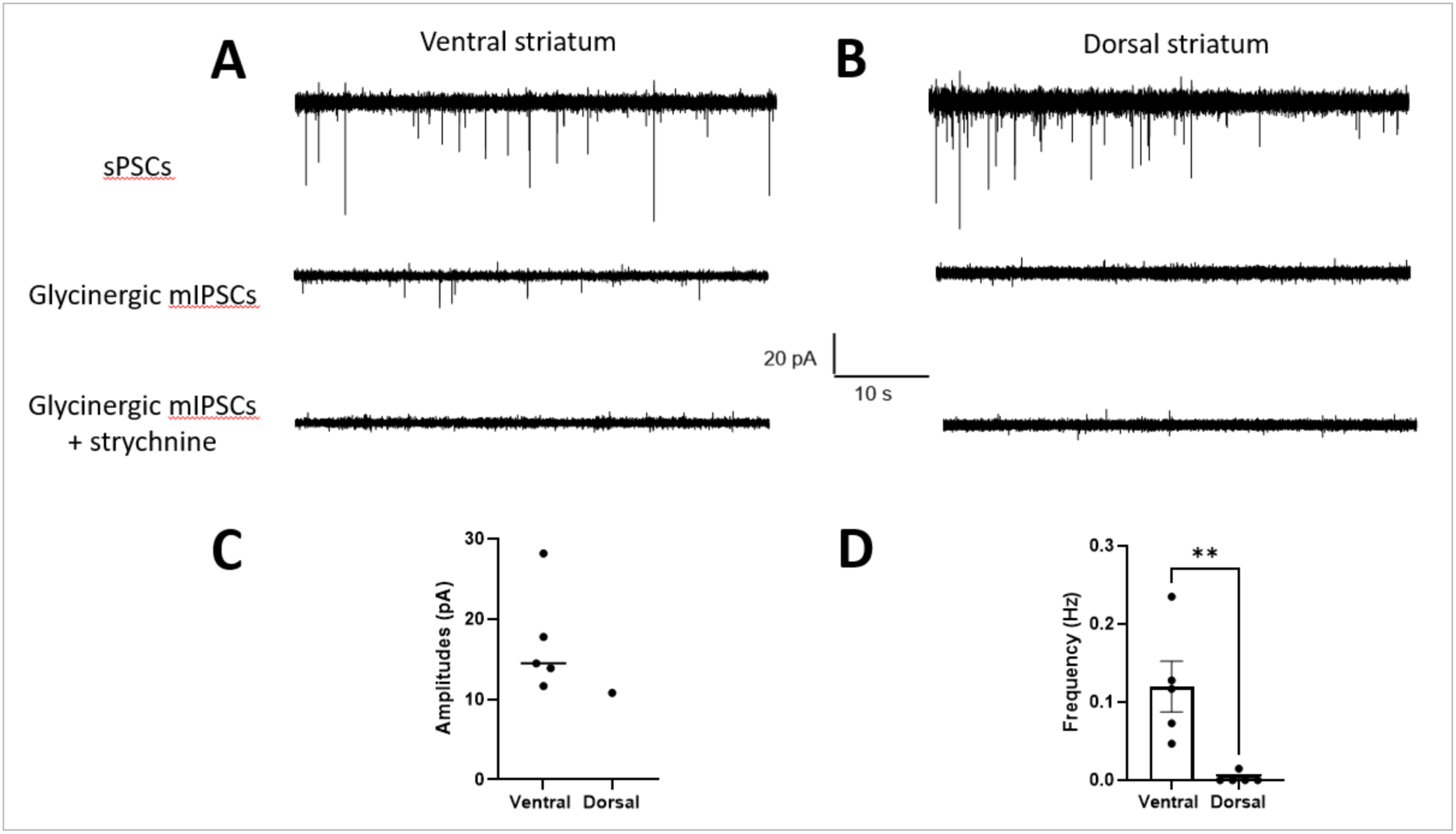
Glycinergic mIPSCs of MSNs in ventral but not in dorsal striatum. **(A-B)** Representative ventral and dorsal current traces recorded using whole cell patch clamp in medium spiny neurons (MSNs) in the ventral (A) and dorsal striatum (B). The top traces show spontaneous postsynaptic current (sPSCs) recorded during aCSF application to confirm whole cell recording. The middle traces show the pharmacologically isolated glycinergic mIPSCs during the application of aCSF containing blockers (10 µM DNQX, 0.1 µM DHBE, 5 µM L-689560, 0.5 µM tetradotoxin, and 10 µM bicuculline) present in the ventral striatum and absent in dorsal striatum. Blocking the mIPSCs by 1 µM strychnine confirms their glycinergic identity. **(C)** Quantification of the amplitude and **(D)** frequency of glycinergic mIPSCs in ventral and dorsal MSNs (mean ± SEM; n = 5 cells from 3 animals in ventral striatum and n = 5 cells from 4 animals in dorsal striatum). Levels of significance determined using a one-tailed Mann-Whitney test (** p < 0.05).

### Expression of recombinant GlyR subunits in cultured hippocampal neurons

Synaptic GlyRs are heteropentamers composed of α (α1–α4) and β subunits, and their assembly and synaptic targeting requires the presence of both types of subunit (Alvarez, 2017). Since the β transcript appears to be highly expressed in most neurons including in the hippocampus (Supplementary Fig. S1), we hypothesised that the expression of α subunits could be the limiting factor controlling the number of synaptic GlyRs. We therefore expressed different recombinant GlyR subunits in cultured hippocampal neurons and analysed their accumulation at inhibitory synapses.

Cells were transduced with lentivirus expressing mEos4b-tagged GlyR α1, α2 or β at day *in vitro* 3 (DIV3), fixed at DIV16, and stained with Sylite to identify synaptic gephyrin clusters. We observed co-localisation of mEos4b and Sylite puncta in neurons infected with either of the virus constructs, indicating that some cultured hippocampal neurons express the corresponding endogenous α or β subunits (Fig. 5A). However, synaptic GlyR puncta were not found in all neurons despite using high virus titres. This means that low levels of α and/or β mRNA transcripts in hippocampal neurons can limit the expression of heteropentameric GlyRs. Interestingly, the mEos4b-GlyRα1 construct was expressed at a higher rate than of the other subunits and had a punctate appearance in most neurons (Fig. 5A). The majority of these puncta co-localised with gephyrin, implying the assembly of mixed GlyRs composed of recombinant mEos4b-GlyRα1 and endogenous GlyRβ subunits. We occasionally also observed mEos4b-GlyRα1 puncta that did not co-localise with gephyrin. These are likely clusters of homopentameric receptors in the extrasynaptic membrane, similar to the clustering of GlyRα3L homopentamers reported in HEK 293 cells (Notelaers et al., 2012). In contrast to GlyRα1, expression of GlyRα2 produced a diffuse labelling in most infected neurons with only weak synaptic puncta. The distribution of recombinant GlyRs at synapses and in the extrasynaptic membrane was confirmed by SMLM imaging of the mEos4b-tagged receptor subunits (Fig. 5B). In line with the above observations, we found a mainly clustered distribution of GlyRα1, both at synapses as well as in the extrasynaptic membrane. The GlyRα2 subunit had a large extrasynaptic receptor population, likely composed of diffusely distributed GlyRα2 homopentamers, and only few synaptic heteropentamers. The mEos4b-GlyRβ subunit was visibly expressed only in a few cells, where it had a predominantly synaptic localisation as a result of its direct interaction with gephyrin.

**Figure 5.**
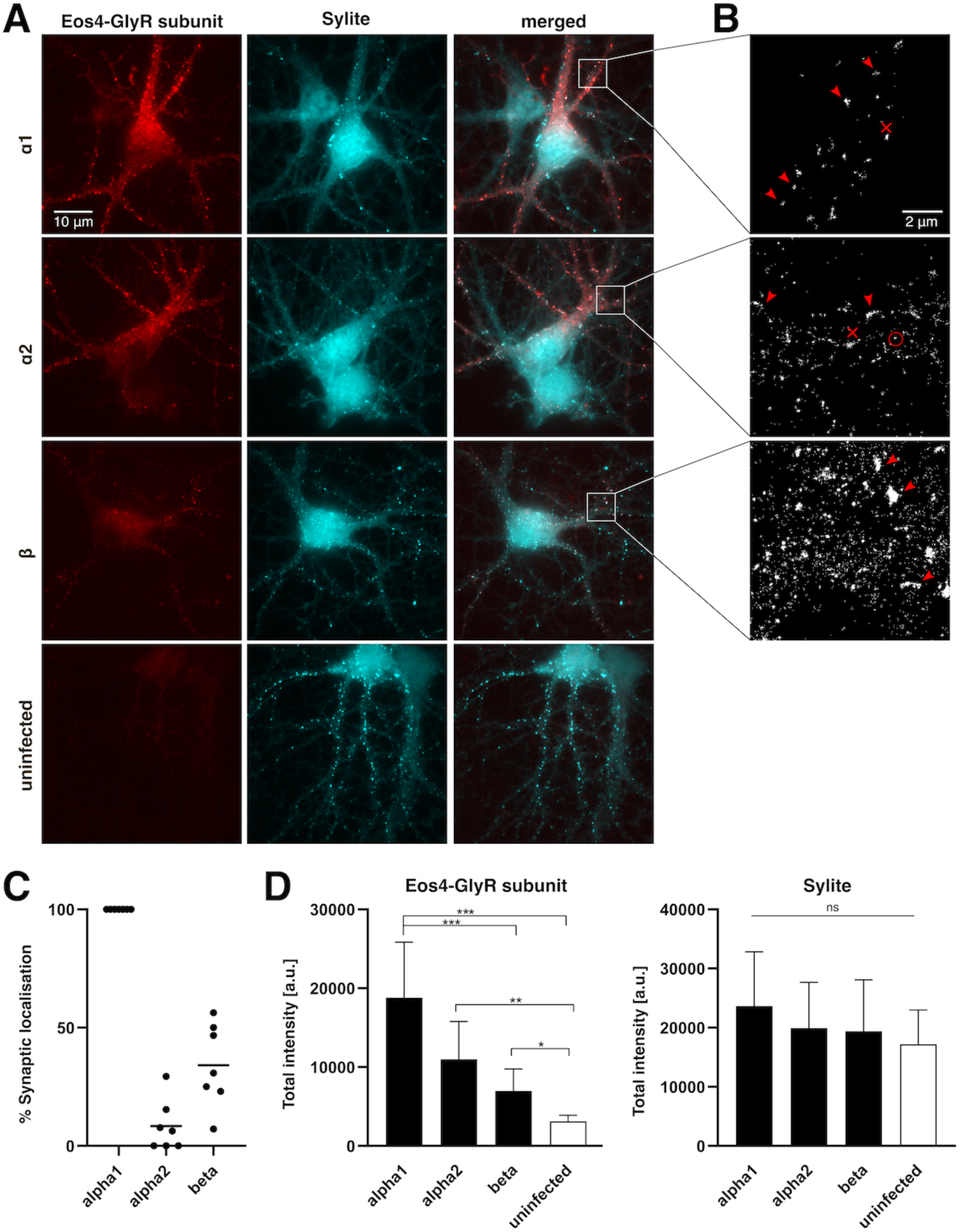
Expression of recombinant GlyR subunits in cultured hippocampal neurons. **(A)** Cultured mouse embryonic hippocampal neurons (E17.5) were transduced with lentivirus expressing mEos4b-tagged GlyR subunits α1, α2 or β (red), fixed at DIV17 and stained for gephyrin (Sylite marker, cyan). Bottom images: uninfected control neurons. Scale: 10 μm. **(B)** SMLM pointillist images showing the photoconverted mEos4b detections. Dense clusters of synaptic (red arrowheads) and extrasynaptic receptors (red crosses) are indicated. Diffusely distributed extrasynaptic GlyR complexes (red circle) are seen as small clusters of detections resulting from the repetitive detection of a single mEos4b fluorophore. Scale: 2μm. **(C)** Quantification of the percentage of infected neurons displaying mEos4b-positive GlyR clusters that co-localise with synaptic gephyrin clusters. Each data point represents one coverslip of cultured neurons (n = 7 coverslips per condition, corresponding to 105 cells for GlyRα1, 114 cells for GlyRα2, and 109 cells for GlyRβ, from N = 4 independent experiments (i.e. cultures). The mean is indicated as horizontal line. **(D)** Quantification of the total fluorescence intensity of mEos4b-tagged GlyR subunits at Sylite puncta in infected neurons and uninfected controls. The integrated mEos4b fluorescence (left graph) and integrated Sylite fluorescence (right) was measured for every Sylite-positive punctum and the median calculated per cell (n = 21 cells for GlyRα1; 9 for GlyRα2; 33 for GlyRβ, and 18 control cells, from N= 3 experiments; mean ± SD; KW test. *p < 0.05; ***p < 0.01; ***p < 0.0001; n.s. not significant). The camera offset was corrected using the minimum pixel intensity in each channel. The signal in the mEos4b channel in the control cultures represents the fluorescence background.

For a more systematic evaluation of the sub-cellular distribution of the GlyRs, we counted the percentage of infected neurons in which a synaptic localisation of the recombinant receptor subunits could be seen in the merged epifluorescence images (Fig. 5C, see Methods). Essentially all GlyRα1 infected neurons had mEos4b-positive gephyrin clusters, indicating a robust synaptic localisation of the α1 subunit. From this can be deduced that most if not all hippocampal neurons express endogenous GlyRβ transcripts. When the cultures were infected with mEos4b-GlyRβ, only few neurons showed a synaptic localisation, meaning that there are only few endogenous α subunits available and that they constitute the limiting factor for synaptic targeting. Interestingly, very few mEos4b-GlyRα2 expressing neurons displayed synaptic localisation, which suggests that not all α subunits are equally capable of forming heteropentameric GlyR complexes with endogenous β subunits.

We also quantified the total intensity of mEos4b-tagged GlyRα1, α2 and β at synaptic gephyrin clusters in infected hippocampal neurons and uninfected control cultures (Fig. 5D). Consistent with our previous observations, mEos4b-GlyRα1 accumulated strongly at inhibitory synapses, again pointing to efficient synaptic targeting. GlyRα2 and GlyRβ subunits showed lesser accumulation at synapses. Pairwise comparison between mEos4b-GlyRα1 and mEos4b-GlyRα2 expressing neurons using a non-parametric Mann-Whitney test showed that the mean total intensity of synaptic GlyRα1 is significantly higher than that of GlyRα2 (p < 0.01, MW test). Although the total intensity of the Sylite puncta was not significantly different between conditions when all groups were compared (Fig. 5E; KW test), a pairwise comparison between mEos4b-GlyRα1 expressing neurons and uninfected controls showed a significant increase in Sylite labelling (p < 0.05, MW test). This seems to indicate that enhanced expression of GlyRα1/β heteropentamers can augment the size of the postsynaptic gephyrin scaffold at inhibitory synapses in the hippocampus.

## Discussion

### Detection of heteropentameric GlyR complexes at hippocampal synapses using SMLM

Making use of the outstanding sensitivity of single molecule localisation microscopy (SMLM) this study describes the presence of very low copy numbers of GlyRs at synapses throughout the hippocampus. Key to this was the use of highly specific labelling of synaptic receptors using a knock-in mouse model expressing endogenous mEos4b-GlyRβ that was used either directly for SMLM through the photoconversion of the mEos4b fluorescent protein or as an antigen that was recognised by a specific anti-mEos nanobody. To positively identify the GlyRβ signals, it was necessary to reduce as much as possible the non-specific background of detections that invariably occurs in SMLM recordings. In the case of mEos4b imaging, this was done by acquiring a negative control movie without photoconversion of the fluorophore. In other words, the photo-physical properties of the fluorophore can be used to identify them with some certainty (Wulffele et al., 2022). Since we focussed on GlyRs at inhibitory synapses, the co-localisation of the detections with the synaptic gephyrin scaffold provided additional assurance that single mEos4b fluorophores could be identified in 10 µm thick slices of brain tissue. These data illustrate the power of SMLM to detect rare or sparsely distributed target molecules in complex samples, in addition to the high spatial precision afforded by super-resolution imaging.

Another strength of SMLM is the possibility to gain access to absolute molecule numbers. Since a fluorophore can be detected several times during the recording, the number of detections has to be translated into the number of emitting fluorophores (Patrizio & Specht, 2016). This is generally more straightforward with fluorescent proteins such as mEos4b that are irreversibly bleached after a number of detections (Wulffele et al., 2022), instead of organic fluorophores (e.g. AF647) that can blink over extended periods. In our case, the use of a knock-in mouse (*Glrb*^eos/eos^) expressing recombinant mEos4b-GlyRβ made it possible to calculate the copy number of endogenous GlyRs at synapses (Maynard et al., 2021). To ensure accurate readings, SMLM movies were recorded until all fluorophores were exhausted (typically 5000 frames in hippocampus and as many as 15000 in the spinal cord, see Methods). Even so, the obtained values have to be taken as estimates given the stochasticity of the number of detections per mEos4b fluorophore.

Our quantification in spinal cord slices indicated that there are on average 120 GlyRs per synapse (Table 1). This result is similar to earlier measurements, in which we calculated median values of 114 and 238 GlyRs at synapses in the dorsal and the ventral horn of the spinal cord, respectively (Maynard et al., 2021). The slightly lower values compared to the earlier study could be related to the lower age of the animals (40 days instead of 2 or 10 months), or to the greater thickness of the sample (10 µm versus 2 µm slices) that affects the signal to noise ratio of the detections. Furthermore, a simpler approach was used in the current study to translate detections into copy numbers. We divided the number of detections at synapses by the number of clusters of detections in sparsely labelled regions in the CA3 that contain mostly individual and spatially separated GlyRs (including extrasynaptic receptors). The advantage of this estimation is that it is independent of the stoichiometry of heteropentameric GlyRs that remains controversial (e.g. (Durisic et al., 2012, Durisic et al., 2014, Grudzinska et al., 2005, Maynard et al., 2021, Patrizio et al., 2017, Yu et al., 2021, Zhu & Gouaux, 2021)). However, our quantification could underestimate the GlyR copy numbers, because a certain fraction of fluorescent proteins like mEos4b is often not functional or not detected under the given imaging conditions, partly due to their complex photo-physical properties (e.g. (Durisic et al., 2014, Patrizio et al., 2017, Wulffele et al., 2022).

As opposed to spinal cord synapses, the numbers of GlyRs at hippocampal synapses are very low. In most cases, only a single cluster of detections (i.e. a single heteropentameric GlyR complex) was present at these synapses (Table 1). Yet, the existing GlyRs appear to be well integrated into the gephyrin domain as judged by dual-colour super-resolution microscopy, pointing to a possible role at synapses (see below).

### Discrepancies between transcription and protein expression of GlyRs in the brain

Our study addresses a long-standing debate about the presence of GlyRs in the brain. According to many reports including recent transcriptomic analyses, GlyRβ mRNA is highly expressed in neurons throughout the telencephalon including the hippocampus (Ceder et al., 2024, Fujita et al., 1991, Malosio et al., 1991) (see also Fig. S1). For instance, ISH data from mouse brain show strong GlyRβ signals in the CA1 neurons consistent with elevated transcription of GlyRβ in pyramidal cells (Allen brain atlas, http://mouse.brain-map.org/gene/show/14434). This is also true for the human brain, where high GlyRβ mRNA levels are detected in all sub-regions of the hippocampus (https://www.proteinatlas.org/ENSG00000109738-GLRB/brain; (Karlsson et al., 2021)). In spite of this, the protein expression of GlyRβ receptor subunits in the hippocampus is very low, as judged by our analysis of synaptic (heteropentameric) GlyRs in mEos4b-GlyRβ knock-in animals. What, then, is the reason for the low protein expression of GlyRβ?

A likely explanation is that the assembly of mature heteropentameric GlyRs depends critically on the co-expression of endogenous GlyR α subunits. The presence of transcripts of the GlyR subunits α1, α2 and α3 in the brain has been demonstrated (Ceder et al., 2024, Malosio et al., 1991), in line with the existence of extrasynaptic (homopentameric) GlyR complexes (Chattipakorn & McMahon, 2002, Molchanova et al., 2017, Mori et al., 2002, Song et al., 2006). Our re-analysis of transcriptomic data confirms that the GlyRα1 mRNA levels are low in the telencephalon, and increase towards dorsal regions of the brain (Fig. S1A), in parallel with the expression of GlyRβ protein and its localisation at inhibitory synapses (Maynard et al., 2021). This raises the interesting possibility that heteropentameric assembly and subsequent synaptic targeting of GlyRs may depend specifically on the concomitant expression of both, α1 and β transcripts. To test this hypothesis, we expressed recombinant GlyR α and β subunits in cultured hippocampal neurons. Lentiviral expression of mEos4b-GlyRβ resulted in synaptic receptors only in some cells, suggesting that in the majority of the neurons the β transcript is not the limiting element for cell-surface delivery and synaptic targeting of heteropentameric GlyR complexes. In contrast, lentiviral infection with an mEos4b-GlyRα1 construct resulted in efficient surface expression of GlyRα1 clusters in hippocampal neurons, where it generally co-localised with the synaptic gephyrin scaffold. In contrast, mEos4b-GlyRα2 was mostly extrasynaptic and less often and less strongly present at synapses. These observations indicate that GlyRα1 (and possibly GlyRα3) could have a particular role in the assembly and forward trafficking of heteropentameric GlyRs towards the plasma membrane.

Our analyses also showed that lentivirus infection did not alter the gephyrin cluster intensities in hippocampal neurons, suggesting that long-term expression of GlyR subunits does not affect the size of inhibitory synapses *per se*. However, the expression of mEos4b-GlyRα1 led to a slight increase in total gephyrin intensity. In our view, this is the result of the increased expression of GlyRα1/β heteropentamers and their accumulation at inhibitory synapses, which in turn can recruit additional gephyrin molecules to the postsynaptic scaffold.

### Possible roles of low-copy GlyRs at brain synapses

The low number of GlyRs at hippocampal synapses begs the question what their role may be. Experiments in GlyRα2 knock-out animals have shown that GlyRs help maintain the excitatory / inhibitory balance in the dorsal striatum (Devoght et al., 2023). However, these receptors are likely to be extrasynaptic GlyRα2 homopentamers. On the other hand, the contribution of a single synaptic GlyR to the chloride influx during inhibitory neurotransmission is probably not significant. In line with this interpretation, glycinergic miniature IPSCs in hippocampal slices are not generally detected (Chattipakorn & McMahon, 2002, Mori et al., 2002, Song et al., 2006), but see (Muller et al., 2013)). The same is true for the *putamen* (dorsal striatum), where whole-cell currents were recorded in response to glycine application (Molchanova et al., 2017). Again, our quantification indicates that GlyR copy numbers in this area are low (Table 1). Synaptic GlyRs are more numerous at synapses in the *nucleus accumbens* (ventral striatum), and indeed, our electrophysiological data support this finding in agreement with earlier studies (Muñoz et al., 2018).

Another possible role of low-copy GlyRs at synapses in the hippocampus, dorsal striatum and maybe other regions of the telencephalon could be a structural one. This concept is based on the high affinity of the GlyRβ-gephyrin interaction (Kasaragod & Schindelin, 2018) that stabilises the receptor as well as the gephyrin scaffold at inhibitory synapses (Chapdelaine et al., 2021). Accordingly, overexpression of synaptic GlyRα1/β heteropentamers led to an increase in gephyrin levels at hippocampal synapses (Fig. 5D). Our two-colour SMLM data of endogenous mEos4b-GlyRβ and gephyrin at hippocampal synapses further confirm that despite their low number the GlyRs are integral components of the postsynaptic gephyrin domain in support of a structural role. GABA_A_R subunits have a much lower affinity for gephyrin and cannot provide the same level of stability (e.g. (Kostrz et al., 2024, Maric et al., 2014, Maric et al., 2011)). However, GABA_A_Rs could probably be recruited efficiently to an existing gephyrin scaffold.

In conclusion, our data positively identify the presence of very small numbers of heteropentameric GlyRs at inhibitory synapses in the brain. This is drastically different to the situation in the *nucleus accumbens*, and even more so in the spinal cord, where GlyRs are abundant and densely clustered at most inhibitory synapses. While it is reasonable to classify different inhibitory synapses as mainly glycinergic or GABAergic, it should be noted that this is a simplification that does not account for the full diversity of inhibitory synapses that may assemble in a continuum of mixed compositions across the entire dynamic range.

## Methods

### Primary culture of hippocampal neurons

Hippocampal neurons were prepared from wild-type mice with Swiss background, at embryonic day E17.5. Experiments were performed according to the European directive on the protection of animals used for scientific purposes (2010/63/EU) and the regulations of the local veterinary authority (Inserm UMS44-Bicêtre, license G94043013). The mice came from excess production for another project, meaning that no animals were generated for the current project.

Pregnant mice were put to death by cervical dislocation and the embryos collected by caesarean section and decapitated. Hippocampi were rapidly dissected in cold Hank’s Balanced Salt Solution (HBSS, Gibco, #14180-046) containing 20 mM HEPES (Gibco, #15630-056) and incubated at 37°C for 15 min in dissection medium containing 0.25% trypsin (Gibco #15090-046). After trypsinisation, hippocampi were washed twice in plating medium composed of Minimal Essential Medium (MEM) containing Earle’s Balanced Salts (EBSS) (Cytiva, #SH30244.01), 2 mM GlutaMAX (Gibco, #35050-038), 1 mM sodium pyruvate (Thermo Fisher Scientific, #11360-039), and 10% heat inactivated horse serum (Gibco, #26050-088). The hippocampal tissue was then triturated in plating medium containing 0.3 mg/ml DNase I (Merck, #11284932001). Neurons were seeded at a concentration of 3.4 x 10^5^/cm^2^ in 12-well plates (Thermo Fisher, #150628) on round glass coverslips (type 1.5, 18 mm diameter; Marienfeld, #0112580) that were pre-coated with poly-D,L-ornithine (Merck, #P8638). The medium was replaced 4 hours after plating with maintenance medium: Neurobasal medium (Gibco, #21103-049) containing B-27 Supplement (Gibco, #17504-044) and 2 mM GlutaMAX. Once a week, 300 µL of fresh maintenance medium was added. Hippocampal neurons were typically infected with ≤ 50 µl of lentivirus stocks at DIV3 (FU-mEos4b-GlyRα2, FU-mEos4b-GlyRβ) or DIV10 (FU-mEos4b-GlyRα1) and fixed for immunocytochemistry at day *in vitro* DIV16.

### Lentivirus expression constructs

The following lentivirus constructs were used for the expression of mEos4b-tagged GlyR subunits: FU-mEos4b-GlyRα1, expressing the coding sequence (cds) of rat *Glra1* isoform a (UniProt P07727-1) (Patrizio et al., 2017); and FU-mEos4b-GlyRβ-bis the cds of human GLRB (UniProt P48167-1) (Grunewald et al., 2018). The receptor sequence in construct FU-mEos4b-GlyRβ-bis (excluding the signal peptide and mEos4b sequence) was replaced with the cds of human GLRA2 (UniProt P23416) to generate the expression construct FU-mEos4b-GlyRα2.

For virus production, HEK-293 tsA201 cells were co-transfected with equal amounts (5 µg each) of the replicon plasmid and the three helper plasmids pMDLg/pRRE, pRSV-Rev, and pMD2.G (Addgene #12251, #12253, #12259) using lipofectamine 2000 (Invitrogen, #11668-019). Cells were cultured in maintenance medium (see above) supplemented with 5 U/ml penicillin and 5 µg/ml streptomycin at 37°C / 5% CO_2_ for 24 h, at which point the medium was exchanged. The culture medium containing lentivirus was collected at 48-55 hours, filtered with a pore size of 0.45 µm, and frozed as aliquots at -70°C.

### Immunocytochemistry in cultured neurons

Hippocampal neurons were fixed at DIV16 with 4% w/v PFA and 1% w/v sucrose in 0.1 M phosphate buffer (PB), pH 7.4 for 10 min. After three washes in PBS, the cells were permeabilized in PBS, pH 7.4 containing 0.25% Triton X-100 and 4% w/v bovine serum albumin (BSA) (Sigma, #A7030) for 10 min and then blocked in PBS containing 4% w/v BSA for 1 h. Neurons were incubated for 1h with the 200 nM of the gephyrin marker Sylite (Khayenko et al., 2022) in PBS containing 1% w/v BSA. The cells were rinsed twice in PBS and kept in PBS overnight at 4°C until imaging.

### Sample preparation of spinal cord and brain slices for SMLM of endogenous mEos4b-GlyRβ

Brain and spinal cord tissue from homozygous mEos4b-GlyRβ knock-in (KI) mice (*Glrb*^eos/eos^, mouse strain C57BL/6N-*Glrb^t^*^m1Ics^, backcrossed into C57BL/6J, accession number MGI:6331106 (Maynard et al., 2021)) was recovered from an earlier project (Wiessler et al., 2024). In this mouse strain the β-subunit of the GlyR is tagged at its N-terminus with the photoconvertible mEos4b fluorescent protein. The chosen animals did not carry any genomic modification other than *Glrb*^eos/eos^. For control experiments brain tissue from a wild-type Swiss mouse was used. Freshly frozen spinal cords and brains from male and female mice at postnatal day 40 were embedded in OCT (Pink Neg-50, Thermo Fisher Scientific, #6502P) and coronal slices of a nominal thickness of 10 µm were cut on a cryostat (Leica, Wetzlar, Germany, #CM3050S) with a chamber temperature of -23°C. The slices were collected on SuperFrost Plus glass slides (Epredia, #J7800AMNZ), fixed with 2% w/v PFA and 0.5% w/v sucrose in PB, pH 7.4 for 10 min and rinsed three times for 5 min in PBS at room temperature (RT). Afterwards, the slices were permeabilized and blocked in PBS containing 4% w/v BSA (A7030, Sigma) and 0.25% v/v Triton X100 for 15 min. Optionally, the slices were labelled overnight with a polyclonal antibody against NeuN (raised in chicken, Synaptic Systems, #266006, 1:2000) at 4°C, followed by 4 h of AF488-conjugated donkey anti chicken secondary antibody (Jackson ImmunoResearch, #703-545-155, 1:2000) at RT. The gephyrin-specific peptide probe Sylite (Khayenko et al., 2022) was applied at a final concentration of 50 nM in PBS containing 1% w/v BSA for 1 h at RT. After two washes of 5 min in PBS, slices were mounted in PBS, covered with a glass coverslip (type 1.5), sealed with PicoDent Twinsil Speed (#1300-1002) and kept overnight at 4°C or directly used for SMLM imaging of the mEos4b photoconvertible protein.

### Single molecule localisation microscopy (SMLM) of mEos4b fluorescent protein

SMLM imaging of mEos4b-tagged GlyRs was performed using an ELYRA PS.1 microscope setup (Zeiss, Jena, Germany). Samples were placed on the motorized stage of an Axio Observer.Z1 SR inverted microscope and imaged with a Plan-Apochromat 63x / NA 1.4 oil immersion objective, with an additional 1.6x lens in the emission path. Images were captured with an Andor iXon 897 back-thinned EMCCD camera (16 bit, 512 x 512 pixels, 16 µm pixel size, QE 90%, set at -60°C working temperature), resulting in an image pixel size of 160 nm. Reference images in the green channel were taken with a 488 nm excitation laser (nominal output 300 mW) and a band pass (BP) 495-575 nm (+ LP 750) emission filter. For the far-red channel we used a 642 nm excitation laser (nominal output 150 mW) and a long pass (LP) 655 nm emission filter. SMLM recordings were performed exploiting the properties of the fluorescent protein mEos4 that is photoconverted from green to red state upon UV illumination (405 nm laser, nominal power 50 mW) and image acquisition in the red channel (561 nm laser, nominal power 200 mW, emission filter BP 570-650 + LP 750). The localisation precision of each detection was calculated from the fitting parameters and was obtained directly from the Zeiss NEO software. For the experiments reported here, the localisation precision was 33.1 ± 12.3 nm in x/y (mean ± SD).

Regarding the experiments in hippocampal cultures (data in Fig. 5), infected neurons expressing mEos4b-tagged GlyR subunits were identified in the green channel, and single reference epifluorescence images of 100 ms exposure were taken with the 488 nm excitation laser set at 1% and the 405 nm laser at 1% of the maximal power, and a camera gain of 100. For Sylite we used the far-red channel, taking one image of 100 ms with the 642 nm laser at 10% output and a camera gain of 10. A SMLM movie of 10000 frames was then recorded at 20 Hz streamed aquisition (50 ms frames) with constant 561 nm laser illumination at 100% laser output corresponding to a maximal power density of 0.95 kW/cm^2^, and a gain 300. The photoconversion of mEos4b-tagged GlyR subunits (from green to red) was done by continuous 405 nm laser illumination that was gradually increased from 0.01 to 4% intensity (≤ 5.3 W/cm^2^ irradiance).

For the detection of endogenous mEos4b-GlyRβ in brain slices (data in Fig. 1 and 3), we used the following acquisition parameters: Hippocampal and striatal regions were identified using NeuN immunolabelling in the green channel (AF488). A single epifluorescence image of 50 ms was taken with 488 nm illumation, 0.2% laser intensity, and a camera gain of 200 to record the NeuN immunofluorescence. Sylite was detected in the far-red channel, taking one image of 100 ms at 642 nm (5% output, camera gain 100). A first SMLM recording of 2000 frames at 20 Hz was done in the red channel with 561 nm laser illumination at 80% (irradiance 0.8 kW/cm^2^), a camera gain of 300, and without 405 nm laser illumination. Under these conditions, mEos4b is not converted into the active (red) form. Afterwards, a second SMLM movie of 5000 frames was taken with the same settings, but with the addition of continuous 405 nm laser illumination that was gradually increased from 0.01 to 3% intensity (irradiance ≤ 4 W/cm^2^). During these recordings, the total population of mEos4b molecules is converted (and bleached), since only very sparse detections are seen at the end of the movie. Glycinergic synapses in spinal cord slices (Supplementary Fig. S2) were identified as bright puncta of (unconverted) mEos4b-GlyRβ subunits in the green channel, and reference images were taken (green, mEos4b: 100 ms acquistion with the 488 laser at 0.2% power, camera gain 200; far-red, Sylite: 100 ms, 642 laser at 3%, gain 200 ms). SMLM movies of 15000 frames were recorded with the same settings as before with a continuous 405 nm laser illumination, gradually increasing from 0.01 to 5% intensity to ensure the complete conversion of mEos4b by the end of the experiment.

### Epifluorescence image analysis in cultured neurons

For the analysis of cultured hippocampal neurons, only transduced cells expressing mEos4b-GlyRα1, α2 or β were considered (data in Fig. 5). Where the expression of the recombinant GlyR subunit was not obvious, this was confirmed by SMLM (Fig. 5B, see below) Within this population, the fraction of cells exhibiting synaptic localisation of GlyR was determined as the number of cells in which mEos4b signals co-localised with Sylite-positive gephyrin clusters, as judged from the epifluorescence images (Fig. 5A). The fluorescence intensity of mEos4b-GlyRα1, α2 and β puncta at synapses was measured only in the infected neurons showing a synaptic localisation of the subunits. Synaptic puncta labelled with Sylite were detected using the spot detector plugin (Icy (de Chaumont et al., 2012)) with the following parameters: bright spots over dark background, detection scale 2 (sensitivity 40), and a size filter between 4 and 100 pixels. To restrict the analyses to synaptic puncta from infected neurons, the signal intensity in the mEos4b channel was thresholded using the mean + 2 SD calculated from uninfected control neurons. The pixel intensity was also corrected for the camera offset, by subtracting the minimum pixel intensity (1500 a.u.) from all images.

### SMLM image analysis of mEos4b fluorescent protein

SMLM movies were processed with Zen software (Zeiss, Zen 2012 SP5 FP3 black, 64 bit) using a peak mask size of 7 pixels, a peak intensity of 6 and excluding the overlapping molecules. The single molecule localisations were corrected for x/y-drift with model-based algorithm (without fiducial markers). A rendered super-resolution image was reconstructed from the pointillist image, in which each detection is represented with a two-dimensional Gaussian distribution with a width corresponding to its point spread function (PSF) and a 10 nm pixel size and saved in tiff format.

Inhibitory synapses in spinal cord and brain slices were identified using the gephyrin-specific peptide marker Sylite (Khayenko et al., 2022). First, the Sylite reference images were adjusted to a 10 nm pixel size by multiplying the pixel number by a factor of 16. We then detected Sylite puncta using the spot detector plugin in Icy with these parameters: bright spots over dark background were detected at scale 5 set to a sensitivity of 100, and the size filtered between 400 and 10000 pixels. The obtained regions of interest (ROI) of Sylite were then employed as mask to measure the intensity of the mEos4b-GlyRβ signals in the rendered SMLM image. Where necessary, the alignment of the reference image and the rendered SMLM image were manually adjusted in ImageJ. To test the specificity of the co-localisation between Sylite and mEos4b-GlyRβ, we performed a pixel shift analysis with images of the CA3 region in Glrb^eos/eos^ slices by horizontally flipping the Sylite channel relative to the mEos4b channel. The number of mEos4b-GlyRβ detections per gephyrin cluster was then recalculated using Icy software. The output data consist in an excel table containing the total intensity of Sylite and mEos4b-GlyRβ in each ROI. To convert the mEos4b-GlyRβ signals into detection numbers, the obtained total intensity values per ROI were divided by the integrated intensity of single mEos4b detections in the image.

A similar approach was used to estimate the copy numbers of mEos4b-GlyRβ containing GlyRs at inhibitory synapses in the CA3 region of the hippocampus. Rendered SMLM images of mEos4b-GlyRβ with a pixel size of 10 nm were segmented in Icy (detection of bright spots over dark background, scale 3 and 4 at a sensitivity of 100, and filtering between a size of 400–10000 pixels). Due to the low mEos4b-GlyRβ density in CA3, the obtained ROIs were considered to be clusters of detections arising from mEos4b fluorophores of a single heteropentameric GlyR (independent of its subunit stoichiometry). The average number of detections per cluster was then calculated by dividing the total intensity of the cluster by the total intensity of a single detection. This value was used as conversion factor to translate detection numbers into molecule numbers and was applied to all SMLM images from spinal cord, hippocampus and striatum. The copy numbers were further corrected by background subtraction using the negative control (CA3 region in wildtype C57BL/6J slices) and are given in Table 1.

### Immunostaining of brain tissue slices used dual-colour SMLM with organic fluorophores

Frozen brain tissue from *Glrb*^eos/eos^ mice (Maynard et al., 2021, Wiessler et al., 2024) was cut into 10 µm thick coronal cryostat sections and processed as described above. For dual-colour SMLM with organic fluorophores, the slices were labelled with primary antibodies against gephyrin (mAb7a mouse monoclonal, Synaptic Systems, #147011, 1:1000 dilution) and mEos protein (AF647-conjugated FluoTag-X2, NanoTag Biotechnologies; #N3102-AF647-L, 1:500) overnight at 4°C, followed by secondary goat anti-mouse antibody coupled with a single CF680 dye (Biotium, #20817, 1:1000) for 4 h at RT. After washing in PBS, the slices were kept overnight at 4°C and imaged the next day.

### Dual-colour SMLM acquisition and image processing

The cover glasses were placed in a closed Ludin chamber (Life Imaging Services) and, through the perfusion holes, was added a homemade dSTORM buffer (Yang & Specht, 2020) that was prepared as follows: A suspension containing 200 μg catalase (from bovine liver, Merck, # C30 1003493507) was washed three times with 1 ml of cold PBS and collected by centrifugation at 12000 g at 4°C for 1 min. After removing the supernatant, the catalase crystals were resuspended in 1 ml of PBS and incubated at 37°C with agitation for 30 min. The final dSTORM buffer was composed of PBS pH7.4, containing 50 mM cysteamine hydrochloride (= β-mercaptoethylamine, MEA, Merck, #M6500), 250 mM glucose (Merck, #G7021), 0.5 mg/ml glucose oxidase (from *Aspergillus niger*, Merck, #G2133-10KU) and 40 μg/ml dissolved catalase. Prior to use, the dSTORM buffer was degassed with N_2_, transferred into a syringe and kept on ice.

SMLM experiments were carried with an Abbelight nanoscope (SAFe 360 Nexus SD) installed on the Zeiss Elyra PS1 setup described above, using two sCMOS cameras (Hamamatsu Orca-Fusion BT) for simultaneous dual-colour imaging. Before the acquisitions, the two cameras were aligned using 0.1 µm TetraSpeck beads (Invitrogen, #T-7279) deposited on a glass coverslip. The Ludin chamber with the tissue slices in dSTORM buffer was then placed on the stage of the inverted microscope and imaged with a Plan-Apochromat 100x / NA 1.46 oil-immersion objective without additional magnification. The far red AF647 and CF680 dyes were excited with a 640 nm laser of 520 mW nominal power (Oxxius LPX-640-500) using adaptable scanning of the excitation region (ASTER). SMLM images were acquired using NEOimaging software v.2.17.1 with the following parameters: 10000 images (image size 256 x 256 pixels and 97 nm pixel size) of 50 ms exposure were taken in the far-red channel with 80% laser power and a field of excitation set at 10%, resulting in a maximal irradiance of 20.3 kW/cm^2^. Fluorophore blinking was gradually adjusted with a 405 nm laser (LBX-405-100, nominal output 108 mW), increasing the power from zero to 2% intensity along the recording (≤ 45 W/cm^2^ irradiance). The emitted wavelengths were separated into two light paths with a 700 nm dichroic mirror for simultaneous dual-colour imaging and filtered with a band pass from 669 nm to 741 nm.

STORM images were processed with NEO analysis software (Abbelight v.39). The raw tiff movies from the transmitted camera (> 700 nm) and the reflected camera (< 700 nm) were loaded and processes sequentially. Single fluorophore signals were detected using temporal mean subtraction with a sliding window of 50 frames, and fit with a Gaussian distribution. The detection coordinates in both channels were corrected for x/y drift and saved as coordinate tables. For spectral demixing and image reconstruction (Supplementary Fig. S4), the detections from the two cameras were imported in the NEO 3D viewer, superimposed and automatically re-aligned. The intensity ratios for each detection were calculated according to the formula I_reflected_ / (I_reflected_ + I_transmitted)_, and we selected for each dye (AF647 and CF680) the inferior and superior cut-offs (0.25-0.32 for AF647); 0.42-0.70 for CF680). After removing the detections that had a precision ≥ 25 nm the spectrally demixed data were saved as coordinate tables and in the form of rendered super-resolution images with a normalized Gaussian representation of each detection and a pixel size of 10 nm. The mean localisation precision was calculated in NEO (Abbelight v.39) and was10.9 ± 4.4 (mean ± SD) for AF647 (anti-mEos nanobody) and 12.2 ± 4.9 for CF680 (anti-gephyrin).

The sub-synaptic distribution of GlyRs (AF647-labelled mEos4b-GlyRβ) and gephyrin (mAb7a-CF680) in hippocampal synapses in the CA3 region was investigated by DBSCAN cluster analysis implemented in NEO 3D viewer (Khayenko et al., 2022), using the following parameters: a radius of ε = 200 nm and n ≥ 5 neighbours for the AF647 detections (GlyRβ); and ε = 80 nm and n ≥ 50 neighbours for CF680 (gephyrin). We exported the data table containing the filtered clusters and their coordinates, and calculated the Euclidian distance between the centre of mass (CM) of the corresponding GlyR and gephyrin clusters, as well as the ratio of the GlyRβ-gephyrin distances divided by the radius of gyration (RG) of the gephyrin cluster. Values <1 indicate that the GlyRβ detections are closer to the CM of gephyrin than the dispersion (RG) of the gephyrin detections themselves, suggesting that the GlyRs are integrated within the postsynaptic gephyrin cluster.

### Confocal microscopy

For the control experiments shown in Fig. S3, spinal cord slices from *Glrb*^eos/eos^ and wildtype animals (*Glrb*^WT/WT^) were labelled with AF647-conjugated nanobody (1:1000, NanoTag) and with an antibody against gephyrin (mouse anti-gephyrin mAb7a, Synaptic Systems, 1:1000), followed by a donkey anti-mouse secondary antibody coupled with CF568 (Sigma #SAB4600075, 1:1000). Confocal images were taken on an SP8 microscope (Leica) using a 63x oil immersion objective and a Hybrid detector (HyD3). Acquisition of images (512 x 512 px, 16 bit) with a pixel size of 120 nm in the x/y plane and 1.27 µm in z (pinhole 1) was done in the three channels, green (mEos4b), red (CF568) and far-red (AF647).

### Statistical analyses

Statistical analyses and graphing were performed using GraphPad Prism v.9. Data are represented as mean ± SD (standard deviation) or mean ± SEM (standard error of the mean) as indicated. Statistical significance was calculated using a non-parametric one-tailed or two-tailed Mann-Whitney test (MW), or a non-parametric Kruskal-Wallis test (KW, one-way ANOVA) with a post-hoc Dunn’s multiple comparison test.

### Preparation of acute brain slices

Adult C57BL/6J mice (postnatal day 35-41) were anesthetized with isoflurane and decapitated. The brains were quickly removed on ice and glued to the cooled stage of a vibratome (Leica VT 1200S). Coronal brain sections containing both dorsal and ventral striatum (150 µm) were sliced in ice-cold cutting solution (140 mM choline chloride, 26 mM NaHCO_3_, 10 mM glucose, 7 mM MgCl_2_, 2.5 mM KCl, 1.25 mM NaH_2_PO_4_, 0.5 mM CaCl_2_, saturated with 95% O_2_ and 5% CO_2_). Brain slices were placed in recovery solution (120 mM NaCl, 2.5 mM KCl, 2 mM CaCl_2_, 2 mM MgCl_2_, 10 mM glucose, 26 mM NaHCO_3_, 1.2 mM NaH_2_PO_4_, saturated with 95% O_2_ and 5% CO_2_) for 45 minutes at 32°C. Slices were kept at 32°C and used 1-4 h after slicing.

### Electrophysiology

Acute brain slices were placed in the recording chamber and perfused with oxygenated aCSF (124 mM NaCl, 4.5 mM KCl, 2 mM CaCl_2_, 10 mM glucose, 1 mM MgCl_2_, 26 mM NaHCO_3_, 1.2 mM NaH_2_PO_4_, saturated with 95% O_2_ and 5% CO_2_, pH 7.4, at room temperature). MSNs in the dorsal or ventral striatum were visualized using 4x air and 40x water immersion objectives of an upright Zeiss microscope (Axioskop 2FS Plus). A P-1000 micropipette puller (Sutter Instruments) was used to prepare filament-containing borosilicate glass patch pipettes (Hilgenberg GmbH) with a resistance of 4-5 MΩ. Patch pipettes were filled with internal solution (120 mM KCl, 4 mM MgCl_2_, 10 mM HEPES, 1 mM EGTA, 5 mM Lidocaine N-ethyl bromide, 0.5 mM Na_2_GTP, 2 mM Na_2_ATP, adjusted to 280 mOsm, pH 7.4). Cells were recorded using voltage-clamp at a holding potential of -60 mV in whole cell configuration with a sampling rate of 10 kHz. During recordings, aCSF bath perfusion was applied for 1 minute. To pharmacologically isolate glycinergic mIPSCs, blockers of AMPA receptors (DNQX, 10 µM), nicotinic receptors (DHBE, 0.1 µM), NMDA receptors (L-689560, 5 µM), action potential firing (tetradotoxin, 0.5 µM) and GABA_A_Rs (bicuculline, 10 µM) were added to aCSF and recorded for 5 minutes. To confirm that these mIPSCs were indeed glycinergic currents, strychnine was added at a concentration of 1 µM to the cocktail of blockers and recorded for an additional 1 minute. All recordings were acquired using a Multiclamp 700B amplifier (Axon Instruments), stored using 1440A Digidata (Axon Instruments), and analysed using Clampfit 10.7.0.3 (Axon Instruments) and NeuroExpress 24.c.16 (Szücs, 2022).

## Data availability

The datasets generated for the current study are available from the corresponding author on reasonable request and will be deposited in a data repository upon article acceptance.

## Acknowledgements

We thank Hans Maric (Virchow Center, University of Würzburg) for the Sylite probe, Carmen Villmann and Natascha Schäfer (Universitätsklinikum Würzburg) for the collection of brain and spinal cord tissue, Krishna Gaete Riquelme (Universidad de Concepción, Chile) for technical assistance, Nicolas Froger (MAPREG, France) for the supply of mouse embryos, Elisabeth Piccart and Petra Bex (UHasselt, Belgium) for brain slice patch clamp training, and Fabrice Schmitt (Zeiss) and Michael Schumacher (NeuroBicêtre, Inserm, UPSaclay) for their generous help with the setting up of the super-resolution imaging platform. We also acknowledge the proficient technical support by Zeiss and Abbelight.

## Author contributions

Experimentation: SC, YV

Data analysis: SC, YV, SYA

Experimental tools and techniques: CM, BB

Project design: SC, BB, CSP

Writing of the manuscript: SC, CSP

Editing of the manuscript: YV, SYA, BB

Funding acquisition: BB, CSP

