## Supplemental information for "Single molecule counting detects low-copy glycine receptors in hippocampal and striatal synapses"

Contents

- Supplementary methods
- Supplementary figures S1 – S4

Supplementary methods

### Single-cell transcriptomic analysis

Data availability: The Allen Mouse Brain transcriptomic data utilized in this study are available through the Gene Expression Omnibus (GEO) database under accession code GSE246717. Specifically, 10x single-cell RNA sequencing (scRNA-seq) data of four datasets, each representing a distinct brain region, ventral striatum (STRv), dorsal striatum (STRd), hippocampus (HIP-CA), and medulla, were accessed through the Sequence Read Archive (SRA) under their run accession codes SRR26528931, SRR26528889, SRR26528942, and SRR26528896, respectively, and retrieved in FASTQ format using the SRA Toolkit (v2.11.0).

Data pre-processing and quality control: Sequencing data were aligned to the mouse genome (mm10, 10x Genomics version 2020-A) using CellRanger (v7.1.0) with default parameters. The filtered gene expression count matrices were then analysed in R (v4.4.1) using the Seurat package (v5.0.1). Each dataset underwent individual quality control (QC) to remove low-quality data and doublets. Briefly, cells were filtered out if they contained <1000 expressed genes or >10% of transcripts derived from mitochondrial genes. Any gene that appeared in <10 cells was removed. Doublets were removed using DoubletFinder package. From around 35 000 cells, approximately 26 000 cells passed QC.

Integration and cell type annotation: The merged Seurat object was normalised using the SCTransform function, with the vst method set to v1 and the rest of parameters set to default, followed by linear dimensionality reduction with PCA. To determine how many principal components to use in downstream analysis, two criteria were used: (1) the cumulative percent variation explained was >90% and the individual percent variation explained was <5%; (2) The change in percent variation was >0.1%. Data integration was performed using the RPCA method, which was selected to account for batch effects across datasets from different brain regions. Further dimensionality reduction and clustering were performed in accordance with the standard Seurat workflow and with the following parameters: UMAP visualisation was performed on the integrated data using the 30 most significant PCs. FindNeighbors was applied on the 2D UMAP embedding (using "umap.rpca" reduction) instead of PCA (default), followed by clustering with FindClusters at a resolution of 0.05, selected to yield a small number of broad clusters corresponding to major mouse brain cell types. Cell-type identities were assigned to the resulting clusters based on the expression of canonical marker genes for neurons (*Snap25, Rbfox3, Syp, Snhg11*), oligodendrocytes (*Mobp, Mog, Plp1, Mag, Mbp*), oligodendrocyte precursor cells (OPC; *Vcan, Pdgfra, Cspg4, Gpr17*), astrocytes (*Gfap, Aldh1l1, Fgfr3, Col23a, Aqp4, Slc1a2, Trp63, Slc7a10, Atp1a2, Gja1*) and microglia (*Ctss, Csf1r, Ptprc, Itgam, Ly86, Myo1f*) drawn from the literature.

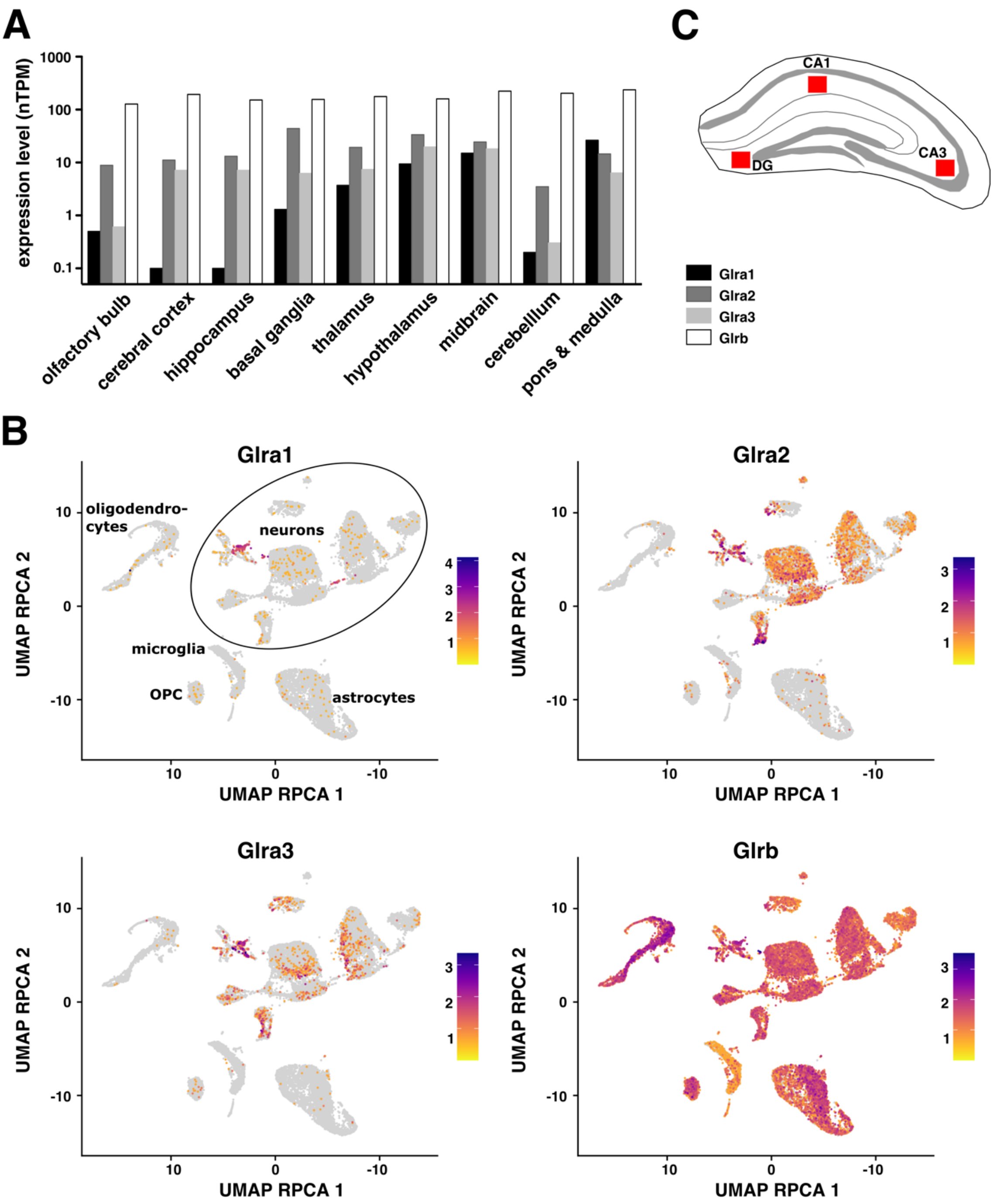


**Figure S1. GlyR gene expression in mouse brain.**

**(A)** mRNA expression of GlyR subunits α1, α2, α3 and β in different brain regions of two month old C57BL/6J mice (sorted from frontal regions on the left to dorsal on the right). RNA-seq data were retrieved from Human Protein Atlas (HPA, https://www.proteinatlas.org) and are expressed as normalised transcripts per million (nTPM). The hippocampus includes the subregions of the *cornu ammonis* (CA) and the dentate gyrus (DG).

**(B)** UMAP projection (using RPCA reduction) of single-cell transcriptomic data derived from four mouse brain regions: hippocampus (HIP-CA), dorsal striatum (STRd), ventral striatum (STRv), and medulla (see Supplementary methods for details on data acquisition and processing). Each panel shows the expression of a different GlyR subunit (Glra1, Glra2, Glra3, or Glrb), visualised with a colour scale from low (yellow) to high (purple). Clusters corresponding to major cell types (neurons, oligodendrocytes, OPCs, astrocytes, and microglia) are labelled in the first panel.
**(C)** Schematic diagram of hippocampus. The red squares represent the hippocampal sub-regions in which SMLM recordings were taken; the molecular layer of the DG and the *stratum radiatum* of CA3 and CA1.

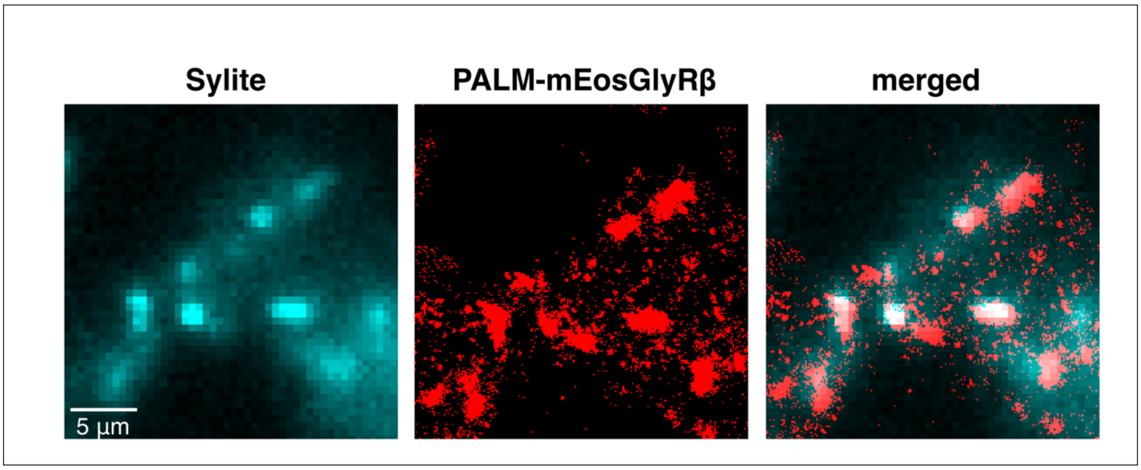


**Figure S2. SMLM of endogenous mEos4b-GlyRβ in spinal cord.**
Single molecule detections of mEos4b-GlyRβ (in red) were recorded in spinal cord slices of *Glrb*$^{eos/eos}$ mice at postnatal day 40 (n = 9733 clusters, 10 fields of view, 3 mice) and labelled with Sylite gephyrin marker (cyan). Scale: 5 µm.

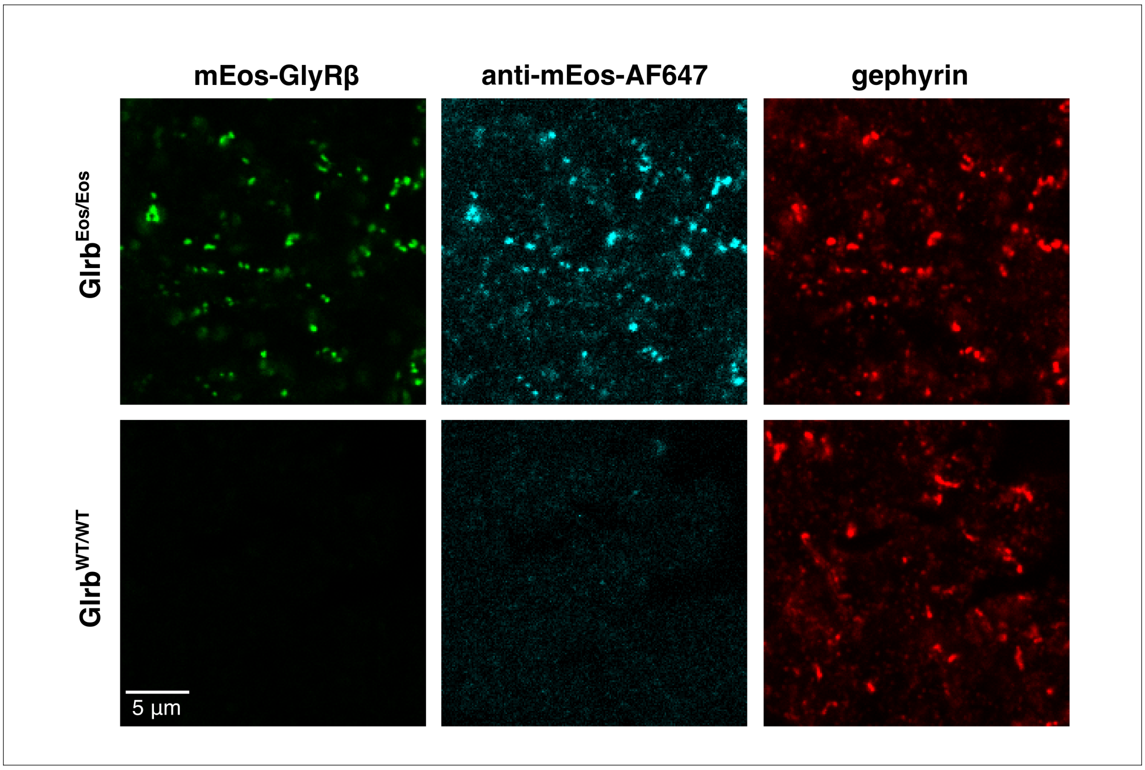


**Figure S3. Specificity of the anti-mEos-AF647 nanobody.**
Spinal cord slices from *Glrb*$^{eos/eos}$ knock-in and wildtype mice (*Glrb*$^{WT/WT}$) were labelled with anti-mEos-AF647 nanobody (AF647-conjugated FluoTag-X2, NanoTag Biotechnologies; #N3102-AF647-L) and with antibodies against gephyrin (mouse anti-gephyrin mAb7a, Synaptic Systems, secondary anti-mouse IgG coupled with CF568). Both the mEos4b-GlyRβ (green) and the anti-mEos-AF647 signals (cyan) co-localised closely with the synaptic gephyrin clusters in *Glrb*$^{eos/eos}$ slices. mEos immunoreactivity was absent in *Glrb*$^{WT/WT}$, showing only minimal non-specific background in conventional fluorescence microscope images. Scale: 5 µm.

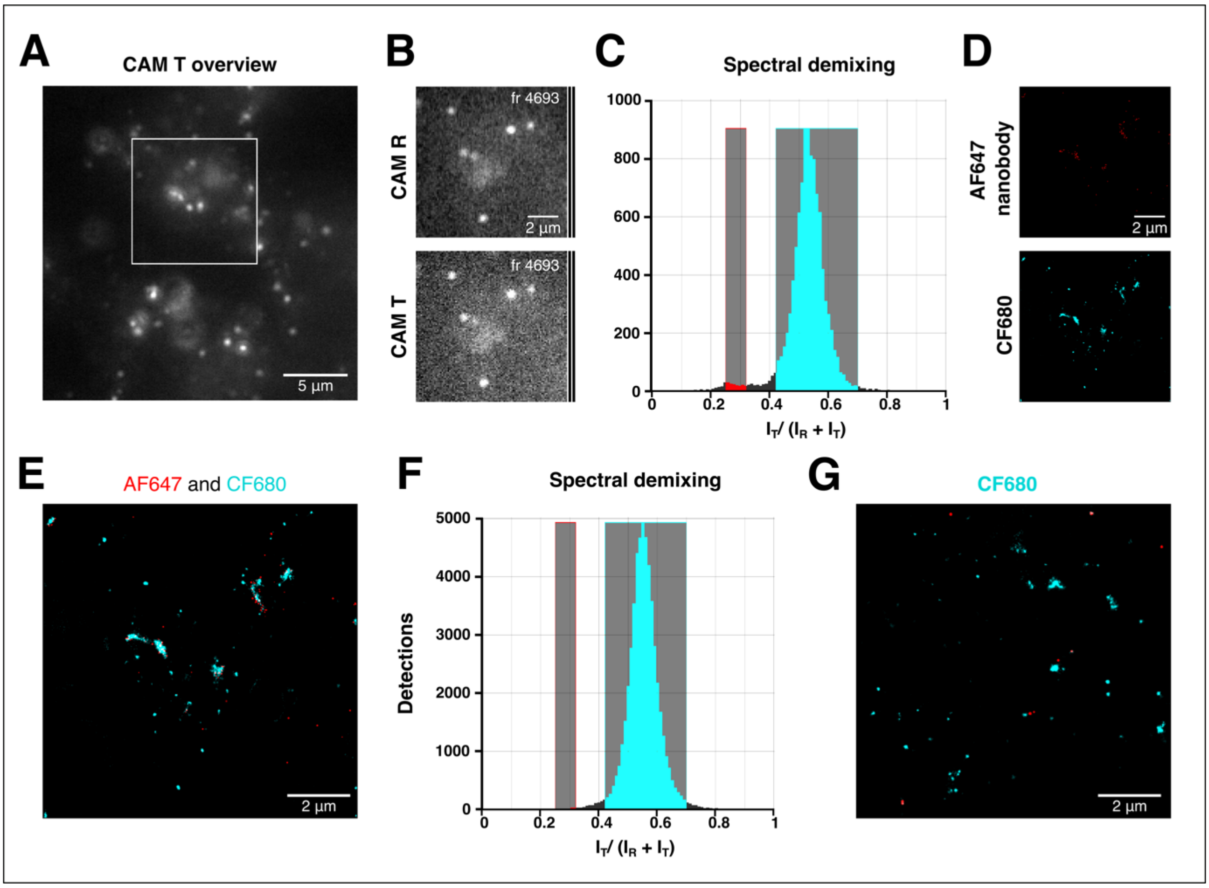


**Figure S4. Dual-colour SMLM using spectral demixing.**
**(A)** Overview from the transmitted camera (CAM T) of endogenous mEos4b-GlyRβ (labelled with anti-mEos-AF647 nanobody) and gephyrin (labelled with mAb7a and CF680-conjugated anti-mouse antibodies) in hippocampal slices of the *Glrb*$^{eos/eos}$ knock-in mouse line at postnatal day 40. Scale: 5 µm. **(B)** Single detections from the reflected (CAM R) and transmitted camera (CAM T) in frame (fr) 4693. Scale: 2 µm. **(C)** AF647 (red) and CF680 (cyan) far-red dyes were separated by spectral demixing, selecting for each fluorophore the inferior and superior cut-offs (0.25-0.32 for AF647; 0.42-0.70 for CF680) of the intensity ratios ($I_{reflected}$ / ($I_{reflected}$ + $I_{transmitted}$)). **(D)** Dual-colour rendered SMLM images of GlyRs labelled with anti-mEos-AF647 nanobodies (red) and gephyrin (cyan, mAb7a-CF680) at hippocampal synapses after spectral demixing. Scale: 2 µm. **(E)** Overlay of the demixed images shown in D. Scale: 2 µm. **(F)** Spectral demixing histogram from a control experiment in which only gephyrin (CF680) was labelled, producing a single peak of intensity ratios. When the same cut-offs as in C were applied, the number of detections in the AF647 channel was significantly reduced (n = 7 recordings, MW test, $p < 0.01$), despite a small number of detections in which the colour was incorrectly attributed. **(G)** Dual-colour rendered SMLM image of gephyrin labelled with mAb7a-CF680 (cyan) in the absence of the AF647-conjugated nanobody. Scale: 2 µm.
